# Somatic mutations in early metazoan genes disrupt regulatory links between unicellular and multicellular genes in cancer

**DOI:** 10.1101/388363

**Authors:** Anna S. Trigos, Richard B. Pearson, Anthony T. Papenfuss, David L. Goode

**Affiliations:** Computational Cancer Biology Program, Peter MacCallum Cancer Centre, Melbourne, VIC 3000, Australia; Sir Peter MacCallum Department of Oncology, The University of Melbourne, Parkville, VIC 3010, Australia; Department of Biochemistry and Molecular Biology, The University of Melbourne, Parkville, VIC 3010, Australia; Department of Biochemistry and Molecular Biology, Monash University, Clayton, VIC 3168, Australia; Bioinformatics Division, The Walter & Eliza Hall Institute of Medical Research, Parkville, VIC 3052, Australia

## Abstract

Extensive transcriptional alterations are observed in cancer, many of which activate core biological processes established in unicellular organisms or suppress differentiation pathways formed in metazoans. Through rigorous, integrative analysis of genomics data from a range of solid tumours, we show many transcriptional changes in tumours are tied to mutations disrupting regulatory interactions between unicellular and multicellular genes within human gene regulatory networks (GRNs). Recurrent point mutations were enriched in regulator genes linking unicellular and multicellular subnetworks, while copy-number alterations affected downstream target genes in distinctly unicellular and multicellular regions of the GRN. Our results depict drivers of tumourigenesis as genes that created key regulatory links during the evolution of early multicellular life, whose dysfunction creates widespread dysregulation of primitive elements of the GRN. Several genes we identified as important in this process were associated with drug response, demonstrating the potential clinical value of our approach.

## Introduction

The spectra of genomic and transcriptomic alterations across cancer types are highly heterogeneous. A multitude of driver genes exist both within and across subtypes (e.g. (Armenia et al., 2018; Nik-Zainal et al., 2016)) spanning a range of mutational types, from simple substitutions, to extensive genome-wide aneuploidies and complex rearrangements (Ciriello et al., 2013; Hoadley et al., 2014; Kandoth et al., 2013; M. S. Lawrence et al., 2013; Zack et al., 2013). This complex array of genetic alterations confounds efforts to assign function to driver genes and identify key driver mutations within patients. Despite this extensive molecular heterogeneity, tumours originating from a variety of tissues converge to several common hallmark cellular phenotypes (Hanahan & Weinberg, 2011). Many of these involve loss of features commonly associated with multicellularity, e.g., uninhibited proliferation, tissue dedifferentiation, disruption of cell-cell adhesion and intercellular communication, suggesting the alteration to genes involved in the evolution of multicellular traits is central to tumour development (Aktipis et al., 2015; Aktipis & Nesse, 2013; Davies & Lineweaver, 2011; Vincent, 2012).

Through the addition of new genes and repurposing of existing genes, metazoan evolution has led to progressive formation of intricate and interconnected regulatory layers (Arenas-Mena, 2017; C. Y. Chen, Ho, Huang, Juan, & Huang, 2014; Schmitz, Zimmer, & Bornberg-Bauer, 2016), which suppress improper activation of replicative processes originating in single-celled organisms and ensure the complex phenotypes and cooperative growth required for multicellularity. Our investigation of gene expression data from a collection of solid tumours revealed extensive downregulation of genes specifying multicellular phenotypes and activation of genes conserved to unicellular organisms (Trigos, Pearson, Papenfuss, & Goode, 2017). This was accompanied by significant loss of coordinated expression of unicellular and multicellular processes within tumours, suggesting selection for the disruption of key regulators mediating communication between unicellular and multicellular genes. Evidence for such selection comes from the clustering of cancer genes at the evolutionary boundary of unicellularity and multicellularity (Domazet-Loso & Tautz, 2010) and the accumulation of mutations in genes required for multicellular development during tumour progression (H. Chen, Lin, Xing, & He, 2015).

Only a limited number of driver mutations are thought to be responsible for the transition from normal, healthy cell to a malignant state (Martincorena et al., 2017; Miller, 1980; Schinzel & Hahn, 2008; Stratton, Campbell, & Futreal, 2009), suggesting individual mutations in highly connected genes in the regulatory network could bring about significant changes in cellular phenotypes. Under our model, the highest impact mutations would be those affecting proteins modulating the communication between the subnetworks that sustain multicellularity and the conserved core of fundamental cellular processes originated in single-celled organisms (Trigos, Pearson, Papenfuss, & Goode, 2018). This would enhance a more primitive phenotype and provide a strong selective advantage for individual cellular lineages. Therefore, the contribution of somatic mutations to the rewiring of transcriptional networks during tumour development can be contextualized and estimated by their effect on gene regulatory networks pieced together during evolution, aiding the identification of key driver mutations.

Here we elucidate how mutational heterogeneity across tumours results in common cellular hallmarks that can be elucidated by accounting for their evolutionary ages and locations in the GRN. We found an overrepresentation of copy-number aberrations and point mutations in genes dating back to early metazoan ancestors. Point mutations disrupted key master regulators that evolved in early in the metazoan lineage, suggesting a primary role in the uncoupling of the subnetworks regulating multicellularity and the fundamental core of cellular processes dating back to single-celled organisms. CNAs were involved in a complementary mechanism of dysregulation, generally disrupting the downstream targets of each of these subnetworks. These results indicate that both point mutations and CNAs contribute to the dysregulation of multicellularity in cancer, but do so in different ways, impacting regions of transcriptional networks with distinct roles in the regulation of multicellularity. Finally, we show how our approach of integration of sequence conservation and transcriptome data with annotated regulatory associations provides a framework for identifying important driver mutations and prioritizing compounds for targeted therapy, at the level of both patient populations and individual tumours.

## Results

### 1. Early metazoan genes are enriched with point mutations and copy-number aberrations acquired during tumourigenesis

We investigated the association between the evolutionary ages of genes and the frequency of copy-number aberrations (CNAs) and point mutations across tumour cohorts. We collected CNA and point mutation data from 3851 and 3867 patients, respectively, from The Cancer Genome Atlas across 7 tumour types (lung adenocarcinoma - LUAD, lung squamous cell carcinoma - LUSC, breast adenocarcinoma - BRCA, prostate adenocarcinoma - PRAD, liver hepatocellular carcinoma - LIHC, colon adenocarcinoma - COAD and stomach adenocarcinoma - STAD). We selected a subset of genes that were consistently amplified or deleted in at least 10% of patients of each tumour cohort, and genes with either missense or loss-of-function (LoF) mutations in at least 3 patients and with a higher rate of occurrence than synonymous mutations (Supplementary figure 1) (*see Methods*). Human genes were classified by their evolutionary age using phylostratigraphy (Domazet-Loso & Tautz, 2010), resulting in 16 phylogenetic groups (phylostrata) (Supplementary figure 2), ranging from genes found in unicellular ancestors (Phylostrata 1-3) (6,719 UC genes), to genes found in early metazoans (Phylostrata 4-9) (7,939 EM genes), and mammal-specific genes (Phylostrata 10-16) (2,660 MM genes) (Supplementary figure 3) (Trigos et al., 2017).

**Figure 1:**
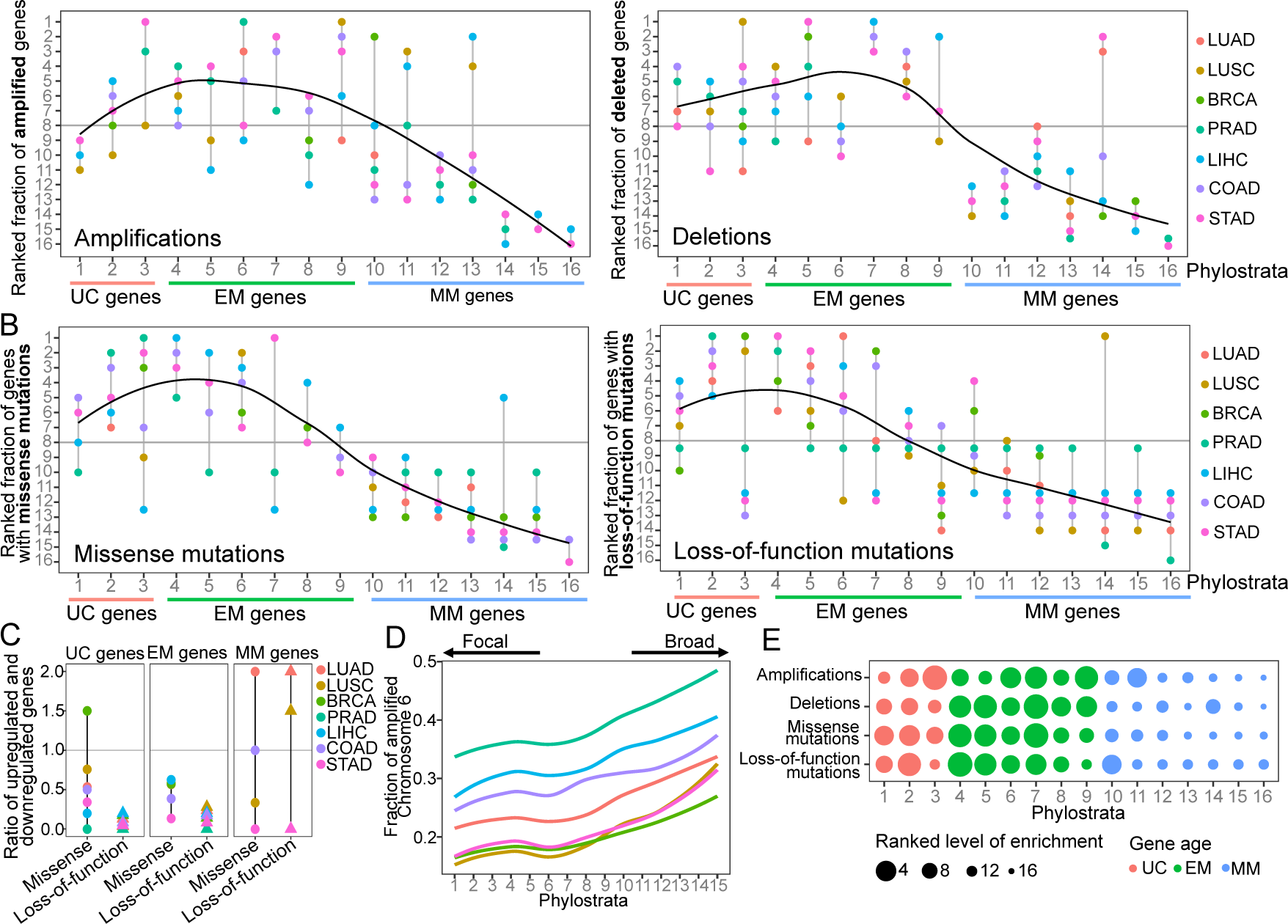
Enrichment of CNAs and point mutations in EM genes. (A) Fraction of amplified (left) and deleted (right) genes across phylostrata. EM genes are preferentially copy-number altered across the 7 tumour types, whereas MM genes are depleted. **(B)** Fraction of genes with missense (left) and LoF (right) mutations across phylostrata. Late UC genes and EM genes are enriched in missense and LoF mutations across tumour types, whereas MM genes consistently have the lowest fraction of genes with point mutations. **(C)** Up and downregulation of genes with point mutations. Ratios greater than 1 indicate a preference for the upregulation of a class of mutated genes, whereas a ratio less than 1 indicates preference for their downregulation. EM genes with missense or LoF mutations are preferentially downregulated across all tumour types. The trend is less evident for UC and MM genes. **(D)** Presence of genes of each phylostratum at different fractions of chromosome altered by amplifications. Chromosome 6 is shown as a representative example. Older genes are preferentially located in regions with focal alterations, whereas younger, MM genes are located in regions with broader changes, suggesting stronger selection for the CNA of UC and EM genes (increasing trend adj. p < 0.05). **(E)** Summary enrichment results of recurrent point mutations and CNAs in phylostrata across tumours. The size of the point corresponds to the level of enrichment (rank). The largest enrichment occurs in EM genes, with some enrichment of UC genes.

**Figure 2:**
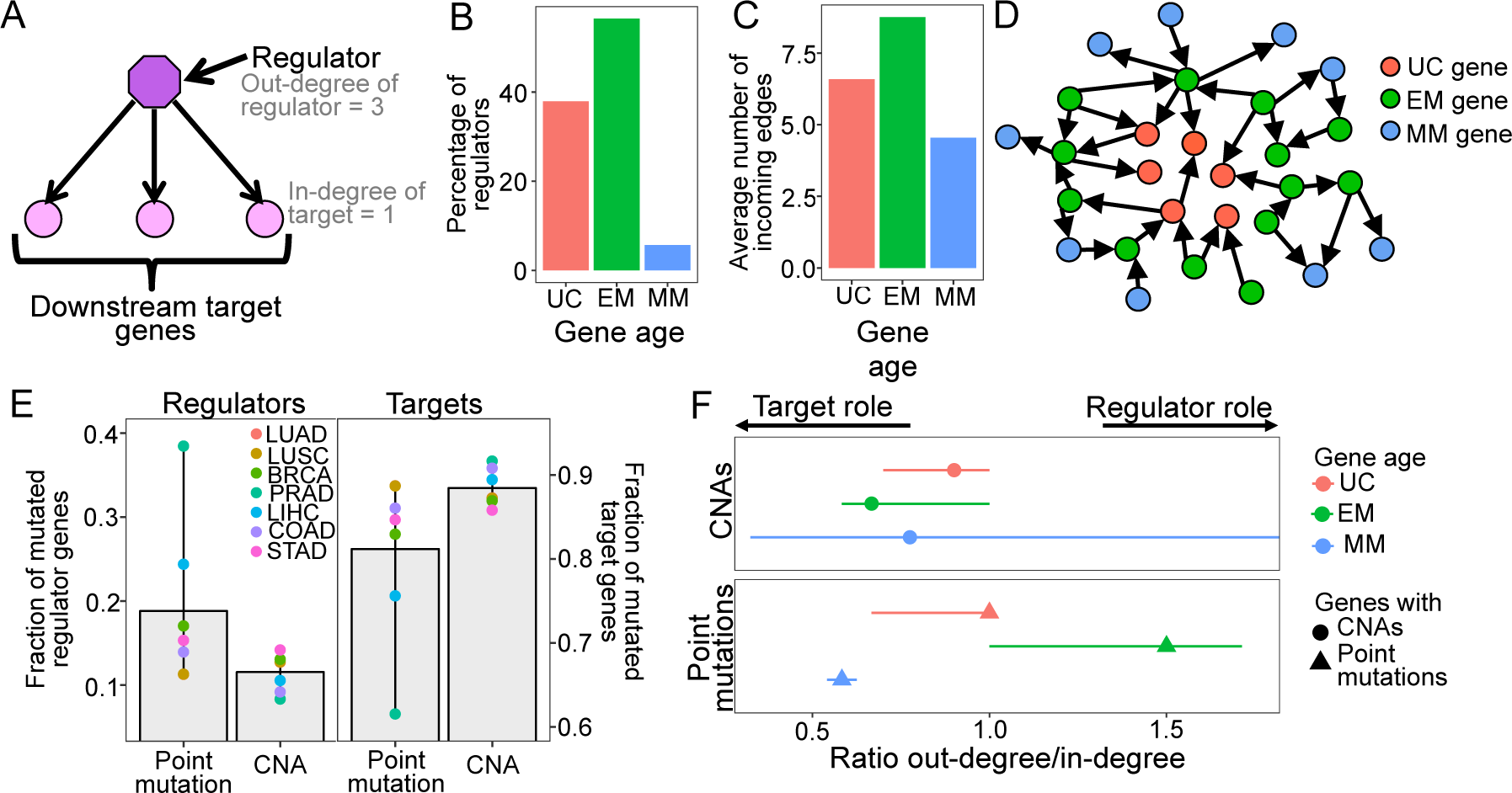
Point mutations in EM genes affect mostly regulators, whereas CNAs in EM genes affect downstream targets. (A) Diagram of a GRN distinguishing regulator and target genes. The number of outgoing edges from a regulator corresponds to its out-degree, whereas the number of incoming edges to a target gene is denoted by its in-degree. **(B)** Percentage of regulators of each age. Regulators are enriched in early metazoan genes (Fisher enrichment test p = 6.48×10^−6^), with over half being EM (56.42%). **(C)** Average number of incoming edges for targets of each age. EM genes are also among the mostly highly regulated genes, with an average of 8.76 regulators controlling their activity, compared to 6.59 and 4.54 regulating UC and MM downstream target genes. **(D)** GRN diagram. EM genes (green) are highly interconnected, acting as master regulators and highly regulated targets. **(E)** Fraction of mutated regulator and target genes by each mutation type. A greater proportion of regulators are affected by recurrent point mutations than CNAs (0.19 vs 0.12; left) whereas the opposite trend is observed for targets (Wilcoxon test p= 0.0078). **(F)** Ratio of out-degree/in-degree of genes with mutations. EM genes are strongly biased towards a preferential regulatory role when point mutated, whereas the CNAs of EM genes preferentially occurs in those with a strong downstream target role. Points represent the median values across tumours and bars represent the upper and lower quantiles.

**Figure 3:**
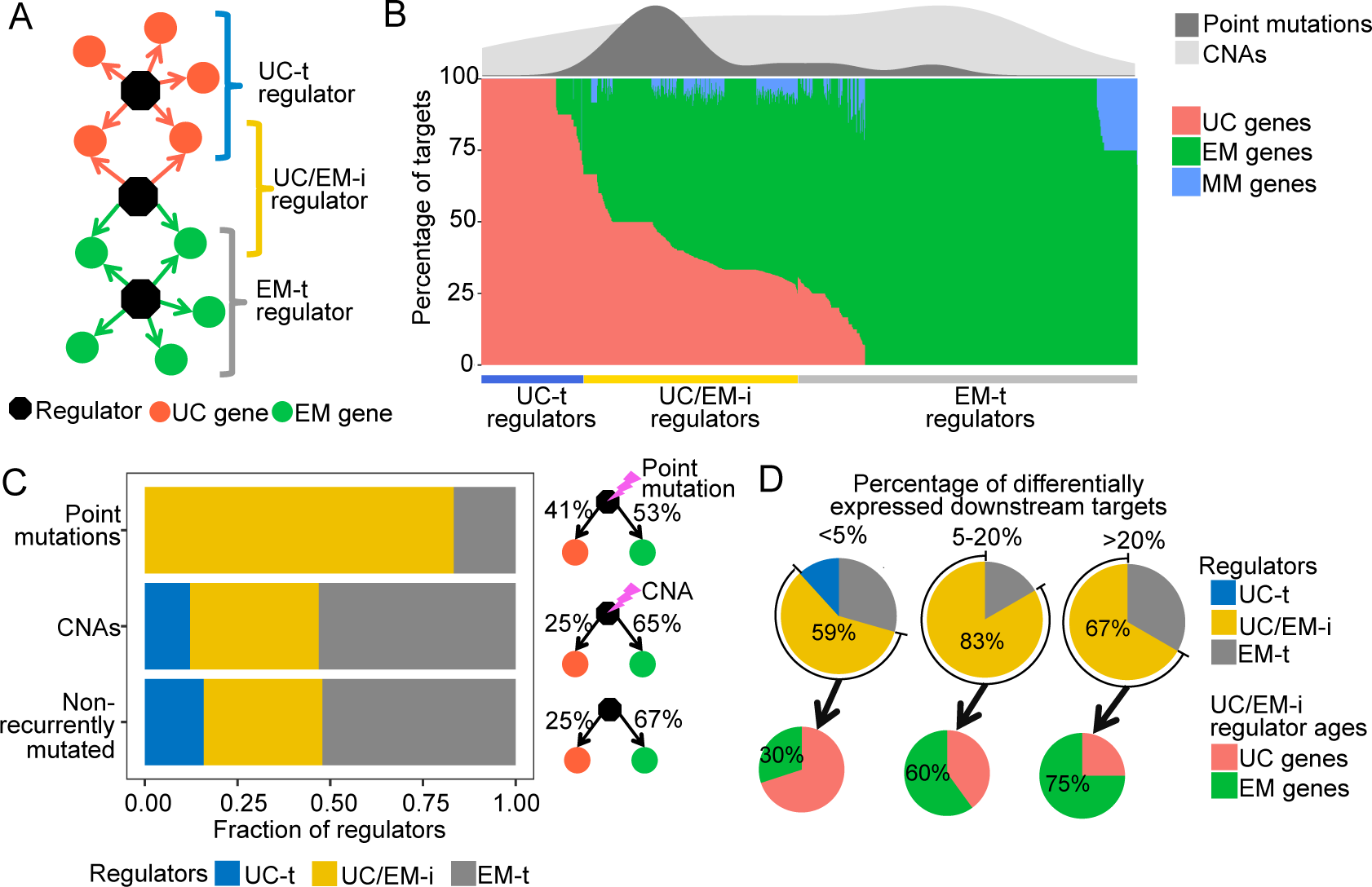
Point mutations in regulators affect UC-EM gene regulation. (A) Classification of regulators by the age of their downstream targets. UC-t regulators mostly regulate UC genes, EM-t regulators EM genes, and UC/EM-i regulators are at the interface of UC and EM genes. **(B)** (Lower panel) Percentage of UC, EM and MM target genes in regulators. (Upper panel) Distribution of recurrent point mutations (dark grey) and CNAs (light grey) across regulators. UC/EM-i regulators are enriched in point mutations. **(C)** Fraction of regulators with point mutations, CNAs and those non-recurrently altered. More than 75% of regulators affected by point mutations are UC/EM-i regulators. The fraction of regulators of each class affected by CNAs is similar to those not affected by recurrent mutations, indicating a lack of preferential alteration of a particular regulator class by CNAs. **(D)** Effect of point mutations in regulators on the expression of downstream targets. Point mutations with a high downstream effect (>20% differentially expressed targets) and a moderate effect (5-20% differentially expressed targets) are more likely to be UC/EM-i regulators of EM origin. Low impact mutations (<5% differentially expressed targets) affect a higher proportion of regulators of a UC origin.

We calculated the fraction of genes in each phylostratum with recurrent CNAs and point mutations, accounting for differences in CNA and mutation rates between tumour cohorts by ranking each phylostratum by the fraction of genes altered (Figure 1A). We found an increasing trend of enrichment of CNAs starting from the earliest UC genes (phylostratum 1), but peaking in EM genes (phylostrata 4-8), with EM genes being the most enriched with both amplifications and deletions across tumours. The majority of tumour types (5/7 tumour types for amplifications and 7/7 tumour types for deletions) had at least 3 EM phylostrata in the top 5 most recurrently altered phylostrata. In contrast, recurrent CNAs were consistently depleted from MM genes (phylostratum 10 onwards), indicating a lack selection for CNAs in younger genes. The decreasing enrichment trend along the phylostrata was significant for amplifications in 5/7 tumour types, and for deletions in all tumour types (Jonckheere-Terpstra tests Benjamini-Hochberg adjusted p < 0.05). Among the recurrently amplified and deleted EM genes are well-known cancer genes. Examples include the EGFR oncogene, recurrently amplified in more than 10% of patients in 5/7 of the studied tumour types and having emerged together with bilaterians (Phylostratum 6), and the tumour suppressor TP53 which also dates back to early metazoan ancestors (Phylostratum 5) and is found recurrently deleted in an average of 20.60% of patients across all tumour types studied. In contrast to the patterns obtained for genes with recurrent CNAs, genes not recurrently copy-numbered altered (frequency < 0.10 across patients) were depleted of EM genes, whereas MM genes are consistently enriched (Supplementary figure 4). An exception is TNFRSF17 (also known as BCMA, BCM), a mammal-specific gene amplified in >10% of BRCA and PRAD patients, involved in immune system processes, which has classically been associated with lymphomas (Laabi et al., 1992) and with oncogenic properties (Coquery & Erickson, 2012; Zhao et al., 2008). Overall, our results suggest that EM genes are specifically preferentially under selection for recurrent CNAs across patients.

**Figure 4:**
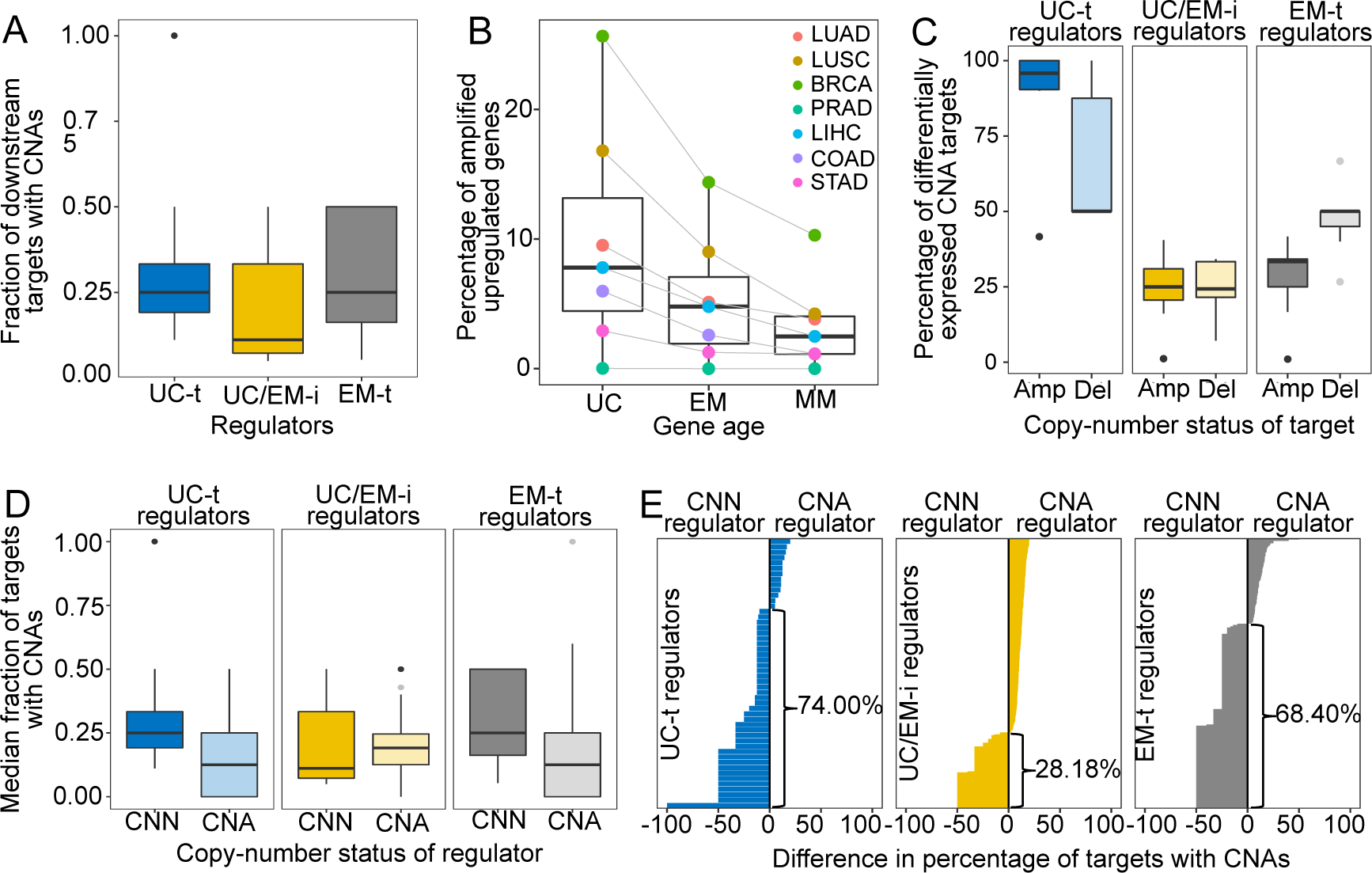
CNAs directly regulate the expression of UC and EM target genes. (A) Fraction of downstream targets with CNAs in regulators. Targets of UC-t and EM-t regulators are more likely to be affected by CNAs than targets of UC/EM-i regulators. **(B)** Percentage of differentially expressed target genes with amplifications. UC and EM target genes are more likely to be upregulated after amplifications compared to younger, mammal-specific genes (Jonckheere-Terpstra decreasing trend test p-value: 0.027). A similar trend is found for the downregulation of deleted genes (Supplementary figure 13). **(C)** Median percentage of differentially expressed CNA genes per regulator class across tumours. Amplified target genes of UC-t regulators are more likely to be upregulated (median 95.83%), than deleted genes are downregualted (50.00%). In contrast, deleted targets of EM-t regulators are more likely to be downregulated (50.00%) than amplifications are upregulated (33.33%). No evident preference is obtained for targets of UC/EM-i regulators (25.00% and 24.29% for amplifications and deletions, respectively). **(D)** Fraction of target genes with CNAs when their regulators are CNA or CNN. A higher fraction of target genes are CNA when UC-t and EM-t regulators are CNN than when they are CNA (Wilcoxon test p = p=6.25×10^−8^ and p=1.76×10^−27^, respectively), indicating a preference for the alteration of targets of these regulators. However, UC/EM-i regulators display the opposite trend, although not significant. **(E)** Difference in the fraction of downstream targets altered by CNAs when their regulators are CNN or CNA. Values less than 0 indicate a higher fraction of CNA targets when the regulator is CNN. This trend is evident across UC-t regulators (74.00%) and EM-t regulators (68.40%), but not for UC/EM-i regulators (28.18%).

A similar enrichment of recurrent mutations in EM genes was identified for point mutations. We found an increasing trend of enrichment of point mutations beginning at genes that date back to later unicellular ancestors (phylostratum 2-3), but peaking in early metazoan genes across tumours, with genes dating to the earliest metazoans (phylostratum 4-5) being the most enriched (Figure 1B). At least 3 of the top 5 most recurrently affected phylostrata by missense mutations were EM in 7/7 tumour types, and by LoF mutations in 4/7 tumour types. In contrast, MM genes were consistently depleted of recurrent point mutations. The decreasing trend of enrichment associated with gene age was significant for missense mutations and LoF mutations in 6/7 tumour types (Jonckheere-Terpstra tests Benjamini-Hochberg adjusted p < 0.05). In contrast, MM genes were harboured the highest fraction of non-recurrently point-mutated genes (Supplementary figure 4).

To determine the effect of somatic mutations on transcriptional states, we compared the expression level of each gene in patients with point mutations and in patients where the gene was not mutated, and calculated the ratio of the number of up and downregulated point-mutated genes of each age in each tumour cohort (Figure 1C). EM genes with missense or LoF mutations were predominantly downregulated, indicating the selection for point mutations in EM genes could be linked to the abrogation of their expression. In contrast, the pattern was less consistent for UC and MM genes, with mixed patterns of preferential up or downregulation according to the tumour and point mutation class considered. These results suggest that the strong selection for point mutations in EM genes across tumours could be linked to their preferential loss of expression.

We also investigated the association between gene age and signatures of selection for CNAs at the chromosome level (Figure 1D, Supplementary figure 5). For each patient and chromosome, we associated the evolutionary ages of the genes located in CNA chromosome regions with the fraction of chromosome affected by CNAs, and found a significant increasing trend in all tumour types (Benjamini-Hochberg adjusted p < 0.005 in all tumour types). UC and EM genes were preferentially located in focally copy-numbered altered regions, suggesting stronger localized selection for the CNA of UC and EM genes. In contrast, MM genes were located in regions of broad copy-number changes, suggesting the CNA of MM genes are likely passenger events swept up in the large chromosomal rearrangements that occur during cancer development.

**Figure 5:**
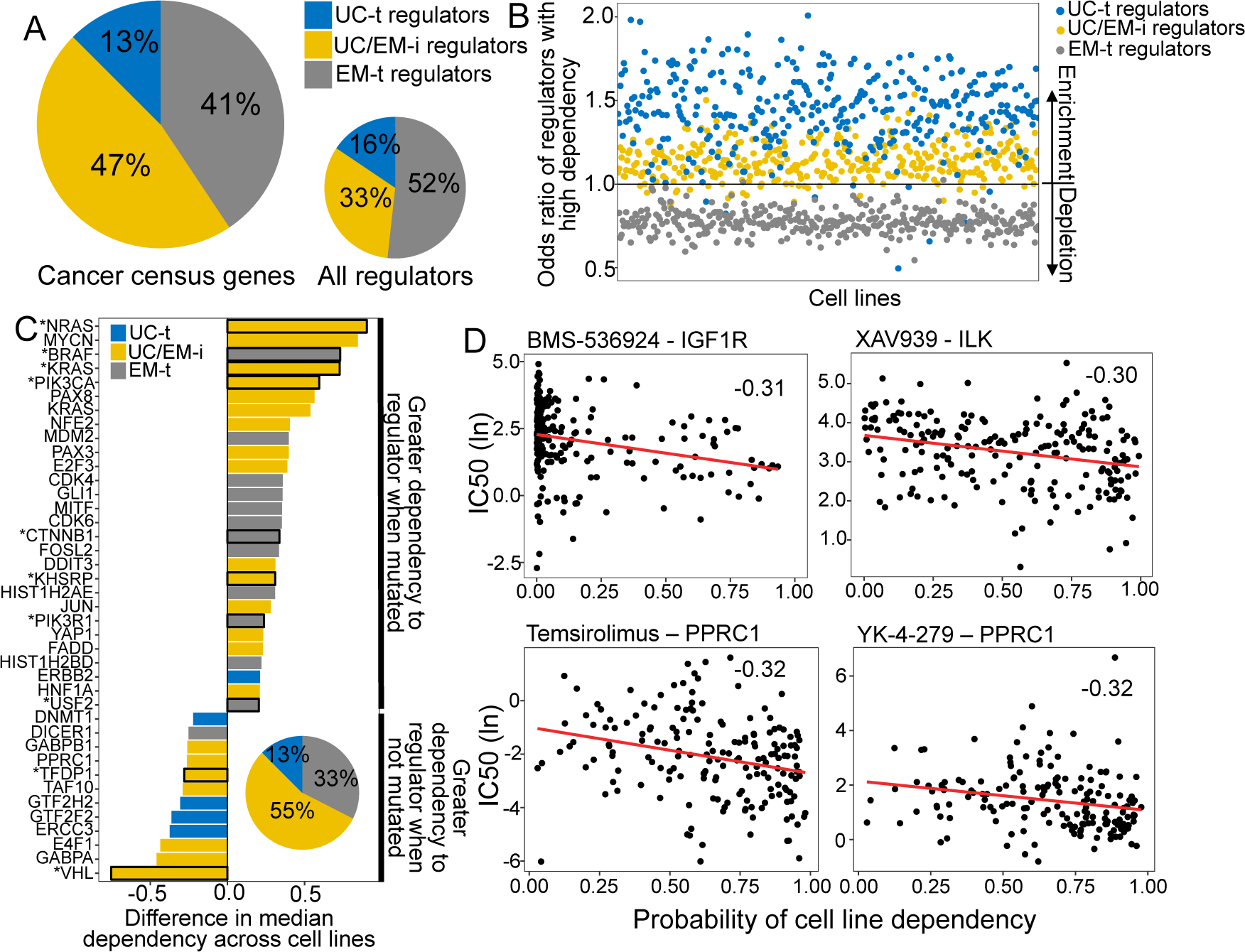
UC/EM-i regulators are fundamental to tumour development and drug response. (A) Fraction of known cancer drivers of each regulator class. While only 33% of regulators are UC/EM-i, 47% of cancer drivers are UC/EM-i regulators, indicating an enrichment of this regulator class in genes involved in cancer development. **(B)** Enrichment of regulators to which cancer cell lines are dependent, as demonstrated by gene knockout. Dependency of cancer cell lines to regulators is associated with regulator class, with an enrichment of UC-t and UC/EM-i regulators and a depletion of EM-t regulators. **(C)** Difference in cell line regulator dependency associated with mutational status. Most cells increase their dependency to specific regulators with point mutations or amplifications, indicating that the mutation of these genes are important for cancer cell survival. This is especially true for UC/EM-i regulators (55%, pie chart). Only regulators with a difference in the median dependency of at least 0.2 are shown. Asterisks denote regulators with significantly different dependency scores between cell lines where the gene was point mutated and non-point-mutated, and the rest those that were significantly different between cell lines where the gene was amplified and non-amplified. **(D)** Correlation between the probability of cell line dependency to UC/EM-i regulators and the IC50 of drugs. (Top left) Expected association between the dependency to the IGF1R regulator and the sensitivity to the IGF1R-inhibitor, BMS-536924. (Top right) Cell lines dependent to ILK show a greater sensitivity to the B-catenin pathway inhibitor (XAV939). (Bottom row) Cell lines dependent on PPRC1 showed increased sensitivity to mTOR inhibitors (temsirolimus) and RNA helicase A inhibitors (YK-4-279).

Our results indicate a preferential recurrent alteration by both CNAs and point mutations of EM genes across tumour types (Figure 1E), suggesting that disruption of these genes by genetic changes likely provides an advantage in the development of multiple tumour types, whereas mutations in genes that evolved later in metazoan evolution, namely MM genes, are unlikely to be playing a significant role.

### 2. Point mutations and CNAs acquired during tumour development differentially affect the human regulatory network

The observed enrichment patterns across cancer types suggested alteration of EM genes provides a selective advantage to tumours. Known cancer drivers are mostly of EM origin (Domazet-Loso & Tautz, 2010) and are highly interconnected in human molecular networks (Cheng et al., 2014), suggesting EM genes hold regulatory roles with important pleiotropic effects in cancer (Trigos et al., 2018). Since important innovations required for the regulation of transcriptional networks from unicellular ancestors evolved in early metazoan species, we investigated whether this could be evidenced in the current structure of the human gene regulatory network (GRN). The GRN was obtained by subsetting the network from PathwayCommons (Cerami et al., 2011) to include only edges annotated with control-of-expression.

Given the directed nature of the GRN, regulator genes can be distinguished from downstream target genes (Figure 2A). As expected, many more genes act as targets (12,812) than as regulators (1,370), indicating the presence of key regulatory hubs regulating a multitude of target genes. We found that over half (56.42%) of regulators in the GRN were EM genes, whereas only 37.88% and 5.69% were UC and MM genes, indicating an enrichment of EM genes as regulators (Fisher enrichment test p = 6.48×10^−6^) (Figure 2B). Focusing on the genes with key regulatory roles, we investigated regulators with at least 10 downstream targets (out-degree >= 10), which correspond to the upperquantile of the distribution of out-degree across all regulators (Supplementary figure 6). We found that 65.12% of these master regulators were EM genes, whereas only 28.49% and 6.40% were UC and MM genes, respectively, indicating that key master regulators with the largest pleiotropic effects in the network were mostly EM genes. This structure of the GRN substantially differed from general protein-protein interaction (PPI) networks (e.g. (Cerami et al., 2011; Chatr-Aryamontri et al., 2017; Li et al., 2017)) where UC genes are usually the most connected (Supplementary figure 7), suggesting specific evolutionary processes shaping the GRN resulted in key regulatory roles for EM genes.

Additionally, EM genes were the most highly regulated downstream genes in the GRN, measured by the number of incoming edges (in-degree), with EM genes having an average of 8.76 incoming edges, compared to only 6.59 and 4.54 in UC and MM downstream target genes (Wilcoxon test p < 2.2×10^−16^ in both cases) (Figure 2C, Supplementary figure 8). The enrichment of EM genes as both regulators and highly regulated downstream targets in the GRN indicates that gene regulation in humans is predominantly under control of EM genes (Figure 2D). Therefore, we hypothesized that the preference of somatic mutations in EM genes might stem from their key regulatory roles in the human GRN.

To test this, we assessed whether selection for somatic mutations in EM genes was linked to their central regulatory roles (Figure 2A). Given that broad CNAs involving large chromosome sections include a high number of genes with poor resolution of the genes under selection, we focused on recurrent CNAs that included less than 25% of the genes of a chromosome in at least 10% of patients of each tumour cohort. We found that point mutations affected a higher fraction of regulators (mean fraction altered = 0.19) than CNAs (0.12) across tumour types (Wilcoxon test p = 0.0078) (Figure 2E), with LoF mutations driving most of the signal (Supplementary figure 9). In contrast, CNAs were more likely to affect downstream target genes without a regulatory role than regulators (Wilcoxon test p = 0.0078) (mean fraction altered = 0.88 for CNAs, 0.81 for point mutations) (Figure 2E). This dichotomy was even more pronounced in somatic mutations that were recurrent in at least 3 of the 7 tumour types. Whereas only 13.89% of genes with recurrent CNAs were regulators, this number was twice as large (26.09%) for genes with point mutations. In contrast, 86.11% of CNAs affected downstream target genes, but only 73.91% of point mutations affected targets. Therefore, recurrent point mutations are more likely to affect master regulators in the GRN, whereas recurrent CNAs preferentially affect genes that predominantly act as targets.

The complex regulatory interactions in the GRN result in many genes having a dual role, acting as both regulators and targets, with EM genes being both master regulators and under high degree of regulation (Figure 2B-D). To account for this dual role, we calculated the ratio of the number of outgoing edges (out-degree) to the number of incoming edges (in-degree), with greater ratios indicating a predominantly regulatory role (Figure 2F, Supplementary table 1), and calculated the median value across all genes with a dual role for each tumour type. We found that for point mutations, EM genes held a predominantly regulatory role across tumours (median ratio = 1.50), whereas UC genes with point mutations did not show this strong bias (median ratio = 1) and MM genes were preferentially downstream genes (median ratio = 0.58). In contrast, EM genes with CNAs were strongly skewed towards being highly regulated downstream targets (median ratio = 0.67) whereas this bias was less marked for UC and MM genes (median ratio UC=0.90 and MM=0.78). Therefore, the observation of selection for point mutations in EM regulators and CNAs in EM target genes also holds for genes with dual regulatory and target roles in the GRN.

Although there is a recurrent selection across tumour cohorts for the somatic mutation of EM genes, our results reveal that selection for point mutations and CNAs differentially disrupt the GRN. EM hub genes with key regulatory roles were preferentially disrupted by point mutations, indicating that few point mutations in key regulators are more likely to create large disruptions across the GRN. In contrast, downstream target genes were preferentially affected by recurrent CNAs, and are therefore more likely to have a localized effect.

### 3. Point mutations disrupt the regulation between UC and EM genes

A main characteristic of cancer development is the loss of coordination in expression between UC and MC genes together with overexpression of UC genes and downregulation of MC genes (Trigos et al., 2017), suggesting a compartmentalization of the GRN into UC and MC gene network regions interconnected by key regulatory links that get disrupted by mutations during cancer development (Trigos et al., 2018) (Figure 3A).

To distinguish these UC and EM network regions, we calculated the percentage of downstream UC and EM target genes for each regulator, and classified individual regulators as preferentially regulating UC targets (>2/3 UC genes) (UC-t regulators), EM targets (>2/3 EM genes) (EM-t regulators), MM targets (>2/3 MM genes) or being at the interface of UC and EM targets by regulating a mix of UC and EM downstream targets (>1/10 UC and EM genes) (UC/EM-i regulators) (Figure 3B, main panel, Supplementary table 2). We excluded regulators that primarily controlled mammalian genes from further analysis, as they only accounted for 2.04% of all regulators. Regulators not meeting any of the above criteria (10.36%) were also excluded. UC-t regulators mostly dated back to UC ancestors (50.81%, Fisher test p =0.021) while both EM-t regulators (56.68%, Fisher test p=0.0011) and UC/EM-i regulators (61.73%, Fisher test p=0.00022) were mostly comprised of EM genes. Given dysregulation of UC and MC gene expression and co-regulation has been found to be consistent across multiple tumours types (Trigos et al., 2017) and therefore likely to share similar drivers, we only considered genes recurrently enriched in point mutations or CNAs in at least 3 of the 7 tumour types studied.

The major constituent of regulators with point mutations were UC/EM-i regulators (83.33%), but these constituted less than half of those altered by CNAs (34.69%) and those not recurrently mutated across tumour cohorts (32.05%) (Figure 3B density plot, 3C main panel). In contrast, only a small subset of point-mutated regulators consisted of EM-t regulators (16.67%) and no recurrently point-mutated gene affected UC-t regulators (Figure 3C). Consistent with a preference for the alteration of regulation between UC and EM genes by point mutations, regulators with recurrent point mutations controlled a similar proportion of UC and EM downstream targets (median target composition of regulators with point mutations = 41.25% UC genes, 53.15% EM genes) (Figure 3C, upper panel). In the case of regulators affected by CNAs, the proportion of UC-t, UC/EM-i and EM-i regulators and the proportions of UC and EM downstream targets was similar to that of non-recurrently mutated regulators (Figure 3C), indicating a lack of selection for CNAs of regulators of a particular age. These results indicate strong overrepresentation of point mutations in regulators at the UC/EM interface, whereas no such trend was observed for CNA regulators or non-recurrently mutated regulators (Supplementary figure 10).

To examine the functional downstream effects of point mutations in regulators, we calculated the percentage of downstream targets that were differentially expressed after point mutations in their regulator genes, and classified the magnitude of the downstream effect as being of low, moderate and high impact according to the percentage of downstream genes affected (<5%, 5-20% and >20%, respectively). The majority of high and moderate impact mutations occurred in UC/EM-i regulators (4/6 and 5/6 regulators) (Figure 2D). Although most low impact mutations also occurred in UC/EM-i regulators (10/17 regulators), these regulators were of UC origin (7/10 of low impact mutations in UC/EM-i regulators), whereas those with high and moderate impact were mostly of EM origin (3/4 and 3/5, respectively). Point-mutated regulators of an early metazoan origin were 2 to 5-times more likely than point-mutated UC regulators to have a moderate or high impact, with 83.33% and 66.67% of regulators with a high and moderate effect being EM genes (Supplementary figure 11). In contrast, point mutations in regulators that resulted in small downstream effects were depleted of EM genes (only 31.82% of genes were EM), indicating that point mutations in regulators that emerged in early metazoan ancestors led to the highest functional impact on downstream targets. Thus, point mutations creating the most substantial alterations to gene expression tend to be in genes of early metazoan origin at the interface of UC and EM genes.

Overall our results suggest that the selection across tumour cohorts of point mutations in EM regulators is mostly tied with transcriptional disturbances of the regulation between UC and EM genes in the GRN, making them potential gene drivers. Multiple known cancer genes were found among UC/EM-i regulators, including PTEN, PIK3CA, MAPK1, MTOR, MYC, NF1, SMAD4, RB1 and TP53BP2. Intriguingly, our analysis pointed to four other EM UC/EM-i regulator genes with mutational frequencies and percentage of differentially expressed downstream targets similar to known cancer genes. Many of these genes, including KEAP1, HNF1A, NFE2L2 and LRRK2, have implied roles in cancer but no strong mechanistic links to date (Figure 5; Discussion).

### 4. CNAs directly regulate the expression of UC and EM downstream targets in the GRN

While somatic mutation of regulators could provide a major selective advantage via simultaneous dysregulation of a multitude of downstream target genes, where and when such mutations occur is largely based on stochastic events during tumour development. An alternate and complementary mechanism for disrupting conserved regions of the GRN without mutation of master regulators would be direct mutation of downstream target genes, as suggested by our finding that CNAs predominantly affected target genes of the GRN (Figure 2E-F).

To investigate the contribution of CNAs to the disruption of the regulatory links between UC and EM genes, we calculated the fraction of downstream targets with CNAs for each regulator in each individual patient. To exclude possible redundant mechanisms resulting from CNAs in both regulators and targets, we only included in the analysis samples where the regulator was copy-number normal (CNN). We found only a small fraction of downstream target genes of UC/EM-i regulators were CNA (median=0.11), whereas a significantly larger fraction of downstream target genes of UC-t and EM-t regulators were affected (median = 0.25 in both cases) (p = 6.14×10^−8^ and p = 4.41×10^−20^ comparing the fraction of CNA targets of UC/EM-i with that of UC-t and EM-t regulators, respectively) (Figure 4A). This indicates that CNAs preferentially affect target genes of UC-t and EM-t regulators, rather than directly disrupting the regulatory links at the interface of UC and EM regions of the GRN.

We hypothesized the preferential of CNAs in targets genes of UC-t and EM-t regulators would be associated with the direct transcriptional modulation of genes by CNAs. To test this, we calculated the expression fold-change in tumour samples with respect to their paired normal samples, and used Wilcoxon tests to compare fold-change values in samples where the gene was CNA and those where the gene was CNN. We found a higher percentage of target genes than regulators were differentially expressed after amplifications across all tumour types, and for deletions in 6/7 tumours types, indicating that CNAs more strongly influence the expression of target genes than regulator genes (Supplementary figure 12). Specifically, UC target genes showed the largest changes in expression after CNAs (median values across tumour types: 7.80% upregulated after amplifications, 8.78% downregulated after deletions), followed by EM genes (4.79% and 5.80%, respectively), and lastly MM genes (2.49% and 3.74%) (Figure 4B, Supplementary figure 13) (Jonckheere-Terpstra decreasing trend test: amplifications p-value: 0.027, deletions p-value: 0.016), indicating that UC and EM targets are more susceptible to changes in expression after CNA.

However, the effect of amplifications and deletions on targets was dependent on regulator class, with a higher percentage of targets of UC-t regulators being differentially expressed after amplifications than deletions increases in expression after amplifications rather than deletions (one-sided Wilcoxon test p=8.29×10^−12^). On the other hand, target genes of EM-t regulators mostly modulated their expression in response to deletions as opposed to amplifications (one-sided Wilcoxon test p=6.09×10^−6^). This trend was also observed when comparing the median percentage of differentially expressed CNA genes per regulator class across tumours (Figure 4C). This suggests selection of CNAs in targets of UC-t and EM-t regulators could be a mechanism for the direct up and down regulation of UC and EM network regions, respectively. In contrast, CNAs of the target genes of UC/EM-i regulators changed their expression much less often, no matter whether it was amplified or deleted (two-sided Wilcoxon test p=0.64), suggesting CNAs are playing a less prominent role in the dysregulation of UC and EM interface regions, where somatic point mutations have a greater impact, but rather directly regulate the expression of UC and EM target genes.

A model of transcriptional changes in UC-t and EM-t targets driven by CNAs as an alternative to direct mutation of the regulators themselves would predict the mutual exclusivity of concurrent CNAs of a regulator and its targets in the same tumour, as the co-occurrence of such mutations would be largely redundant. To test this hypothesis, we calculated the median fraction of targets with CNAs for each regulator across all patients in the 7 tumour cohorts, and found that the fraction of targets with CNAs was significantly higher in patients where the regulator was CNN than when the regulator was CNA (Wilcoxon test p-value = 3.34×10^−20^, Supplementary figure 14). However, this trend was only observed for targets of UC-t and EM-t regulators (one-sided Wilcoxon test p=6.24×10^−^ 8 and p=1.76×10^−27^, respectively), but not for targets UC/EM-i regulators (p=0.77), where there trend seemed to be the opposite (Figure 4D), indicating that preferential alteration of target genes by CNAs, as opposed to regulators, is a mechanism employed for the independent modulation of UC and EM network regions.

We next investigated the fraction of regulators whose targets were preferentially modulated by CNAs. For this, we calculated the difference in the fraction of downstream target genes when regulators were CNN and CNA (Figure 4E). We found that over two thirds of UC-t and EM-t regulators (74.00% and 68.40%, respectively) had a higher fraction of copy-aberrant downstream genes when they were CNN than when they were CNA. These results were independent of the evolutionary ages of the regulators, since the percentages were similar for both UC and EM regulators (Supplementary figure 15). A potential explanation for this pattern is illustrated by MDM2, where 33.33% of its downstream targets were CNA when MDM2 was CNN, but no target was CNA when MDM2 had changed in copy-number. Since MDM2 is an inhibitor of p53, either amplification of MDM2 or deletion of p53 would have a similar effect, and therefore there is no selection for the simultaneous co-occurrence of both alterations in the same patient. In contrast to the strong trends observed for UC-t and EM-t regulators, this trend was only observed in 28.18% of UC/EM-i regulators.

Here we found that CNAs are a widespread mechanism of dysregulation of UC and EM target genes, directly modulating the expression of targets of UC-t and EM-t regulators. The effect of CNAs on the GRN is therefore distinct to the disruption by point mutations of the regulatory links between UC and EM genes, but are also important drivers of large transcriptional disturbances in tumours.

### 5. UC/EM-i regulators are important drivers of tumourigenesis and influence drug sensitivity

UC/EM-i were found to be preferentially targeted by point mutations, and are likely to be key points of vulnerability to cancer development given their regulatory role modulating UC and EM genes (Trigos et al., 2018). Therefore, we investigated these regulators as potential drivers of tumourigenesis and their role in determining drug response.

We compiled a set of known cancer drivers from the Cancer Census COSMIC database (Forbes et al., 2017). These genes were enriched in EM genes (Fisher test p = 0.0015, 57.36% EM genes, 39.58% UC genes and 3.06% MM genes). 36.71% of the known cancer genes were regulators, and were enriched in UC/EM-i regulators (46.88%, Fisher test p= 0.0043), but depleted in UC-t (p=0.83) and EM-t regulators (p=0.95) (Figure 5A), supporting modulation of regulation between UC and EM genes is a common effect of cancer drivers.

To determine the importance of these regulators to cancer development, we used the dependency scores from CRISPR-Cas9 essentiality screens of 364 solid-tumour tissue cell lines from the Avana CRISPR-Cas9 genome-scale knockout dataset made available by Project Achilles and the Cancer Dependency Map project (Meyers et al., 2017). A high probability of dependency indicates a gene is essential for proliferation of a given cell line (Supplementary figure 16) (*for further details, see* (Meyers et al., 2017)). We calculated the odds ratio (OR) of each regulator type having a large effect on cell line proliferation (probability of dependency >= 0.95), with OR greater than 1 indicating increased likelihood of a high dependency (Figure 5B, Supplementary figure 17). As expected, we found most cell lines were highly dependent on UC-t regulators (OR > 1 in 96.98% of cell lines), likely due to their role in regulating fundamental functions required for cell survival. However, we also found that 92.86% cancer cell lines were highly dependent on UC/EM-i regulators (OR consistently greater than 1), indicating these regulators are fundamental for cancer cell survival. In contrast, cancer cell lines were not nearly as dependent on EM-t regulators for their proliferation (OR > 1 in only 0.55% of cell lines), indicating dysregulation of EM processes might contribute to tumourigenesis, but not be sufficient by themselves. These results suggest that UC/EM-i regulators are indispensible for cancer cell line proliferation.

We hypothesized the degree of dependency on UC/EM-i regulators would be tied to their mutational and copy-number status. For each regulator, we classified cell lines into those with or without a point mutation or amplification in the regulator based on data from the Cancer Cell Line Encyclopaedia (CCLE) (https://portals.broadinstitute.org/ccle), and calculated the median dependency scores for mutated and non-mutated cell lines (Supplementary table 3). We only considered regulators whose difference in median dependency between mutated and non-mutated cell lines was at least 0.20, and were significant by Wilcoxon tests (p < 0.05). This revealed 40 regulators whose mutation was associated with cell-line dependency (Figure 5C). Multiple well-known cancer genes were among these top hits. Amplification of the oncogenes ERBB2, CDK6, MDM2 affected dependency, as did point mutations in PIK3CA, KRAS, NRAS, VHL and BRAF, validating our approach to highlight genes with significance in cancer. Over half of these top hits correspond to UC/EM-i regulators (55.00%), indicating that mutations in UC/EM-i regulators are more likely to affect the dependency of a cancer cell line to a regulator. We found that in most cases cell lines were more dependent on a regulator when it was point mutated (80.00%) or amplified (66.67%) than when it was non-mutated, suggesting that mutation of these regulators are key to cancer cell line proliferation (Figure 5C).

We also investigated the implications of dependency to UC/EM-i regulators on drug sensitivity. Based on the half maximal inhibitory concentration values (IC50) from drug sensitivity screens covering 250 drugs from the Genomics of Drug Sensitivity in Cancer database (Yang et al., 2013), we calculated the Spearman correlation between dependency scores and the IC50 values (Supplementary figure 18), and identified 11 significant associations with UC/EM-i regulators, 38 with UC-t regulators and 23 with EM-t regulators (correlation < −0.25 & adj. p < 0.05). These identified regulators were among the most highly correlated with the IC50 of the particular drug (Supplementary figure 19). As expected, we found strong correlation between dependency scores for UC/EM-i regulators and the IC50 of drugs targeting the regulator (e.g. IGF1R and Linsitinib, BMS-536924 and BMS-754807), as well as between MAPK1 and an inhibitor of related genes in the MAPK/ERK pathway ((5Z)-7-Oxozeaenol), validating our approach (Figure 5D, Supplementary figure 20).

However, we also found unexpected strong correlations between the IC50 of particular drugs and the dependency scores of UC/EM-i regulators (Figure 5D, Supplementary figure 20). For example, the IC50 of XAV939, an inhibitor of Wnt/β-catenin, was also strongly correlated with the dependency to ILK (−0.30), a regulator of integrin-mediated signal transduction involved in tumour growth and metastasis, supporting the use of Wnt/β-catenin inhibitors for cancers dependent on ILK, including colon, gastric and ovarian and breast cancers (Hannigan, Troussard, & Dedhar, 2005). We also found strong correlation across cell lines between the dependency to PPRC1 and mTOR-inhibitors (temsirolimus, used in the treatment of renal cancer), dual PI3K/mTOR-inhibitors (dactolisib, in clinical trial for advanced solid tumours (Wise-Draper et al., 2017)), YK-4-279 (showing pre-clinical efficacy for Ewing sarcoma (Lamhamedi-Cherradi et al., 2015)) and the chemotherapy agent docetaxel, currently used in the treatment of breast, lung cancer, stomach cancer, head and neck and prostate cancer. Of the tumour types included in our study, the correlation with PPRC1 dependency was particularly strong (< −0.25) in liver, lung and stomach cell lines for temsirolimus sensitivity, lung and stomach cell lines for docetaxel and dactolisib sensitivity and breast cell lines for YK-4-279 sensitivity, but were also held for a number of other solid tumour types (Supplementary figure 21), suggesting their use across multiple cancer types. With this, our novel approach has identified novel potential vulnerabilities for cancer development and proposes drugs repositioning.

## Discussion

Detailed analyses of recurrent somatic mutations across tumour types revealed the prevalence of mutations related to both gene age and its position within the regulatory network. We provide evidence that point mutations and CNAs play complementary roles in the transcriptional dysregulation in cancer by affecting distinct regions of the underlying gene regulatory network, supporting the loss of communication between the core biological processes originating in ancient single-celled life and the regulatory controls acquired during metazoan evolution to control these processes. This would result in tumour convergence to similar transcriptional states of consistent activation of genes from unicellular ancestors and loss of cellular functions characteristic of multicellular organisms. Our results attribute key roles to genes at the interface of unicellular and multicellular regulation in tumourigenesis, with implications for conventional and experimental therapies.

Common hallmarks shared by tumours of diverse genetic backgrounds suggest the consequences of mutations acquired during tumour development follow common principles, promoting the downregulation of genes and pathways associated with multicellularity and the activation of fundamental cellular processes evolved in early unicellular organisms (Trigos et al., 2017). Here we found genes central to the human gene regulatory network that arose in early metazoans were the most often recurrently affected by point mutations and CNAs across tumour types. Other studies have found that gatekeeper cancer drivers (those that regulate cell cooperation and tissue integrity) emerged at a similar evolutionary time, whereas caretaker genes (those ensuring genome stability) emerged at the onset of unicellular life (Domazet-Loso & Tautz, 2010). Our results suggest recurrent mutations mostly affect gatekeeper genes regulating fundamental aspects of multicellularity, whereas the disruption of caretaker activities by recurrent somatic mutations and CNAs is more limited.

We found the impact of point mutations and copy-number aberrations was concentrated on specific regions of the gene regulatory network. Point mutations preferentially affected gene regulators at the interface of unicellular and early metazoan subnetworks, resulting in a loss of regulation of multicellular components over fundamental cellular processes. On the other hand, CNAs preferentially affected their downstream target genes, directly promoting the independent activation and inactivation of regions predominantly composed of unicellular and multicellular genes, as opposed to mixed regions, further supporting a loss of multicellularity and the tight association between gene expression level and gene age (Trigos et al., 2017).

Samples where UC/EM-i regulators were mutated tended not to have CNAs in the corresponding target genes and vice versa; in samples carrying CNAs for multiple target genes, unicellular and multicellular, the cognate UC/EM-i regulator were mostly unaltered. These complementary but distinct mechanisms of alteration to the same regulatory subnetworks by different classes of somatic mutations in different tumours provide mutually exclusive but highly convergent paths towards common hallmarks associated with the loss of multicellularity features in cancer.

Our evolutionary network analysis approach also highlights potential early driver genes by elucidating the evolutionary regulatory context in which genes operate. Only a handful of driver mutations are thought to be responsible for the transition from a normal, healthy cell to a malignant state (Martincorena et al., 2017; Miller, 1980; Schinzel & Hahn, 2008; Stratton et al., 2009), but their identification amid large and diverse genetic alterations is challenging. Our results suggest the key positioning of early metazoan regulator genes at the interface of network regions from unicellular and multicellular ancestors makes cells susceptible to widespread dysregulation of transcriptional networks upon their disruption, as their alteration would uncouple the regulatory controls required for multicellularity (Greaves, 2015). This implicates these genes in key roles in the onset and progression of cancer and highlights them as potential gene drivers and drug targets. Our analysis of the effect on cell line dependency after knockout of these regulators revealed that their alteration is capable of modulating cell proliferation across 364 cell lines.

Many of these interface regulators have not been significantly studied in the context of cancer, but drugs targeting these genes are currently in clinical trial for other diseases, opening the possibility for drug repurposing. LRRK2 encodes the dardarin protein, considered to be central to the aetiology of Parkinson disease (Zimprich et al., 2004). Two inhibitors of dardarin, DNL201 and DNL151, are currently undergoing testing in clinical trials as a means to slow down or regress neurodegenerative diseases (Atashrazm & Dzamko, 2016). We also identified that cell lines dependent on the UC/EM-i regulator PPRC1, peroxisome proliferator-activated receptor gamma, co-activator-related 1, were particularly susceptible to mTOR inhibitors, YK-4-279 and docetaxel. PPRC1 is an activator of mitochondrial biogenesis, a process regulated by mTOR (Morita et al., 2013; Ramanathan & Schreiber, 2009; Schieke et al., 2006), highlighting the use of mTOR inhibitors in cancers with aberrant mitochondrial activity. Furthermore it suggests that YK-4-279, a binding inhibitor of oncogenic fusion proteins in Ewing’s sarcoma and with encouraging pre-clinical efficiency in this cancer (Lamhamedi-Cherradi et al., 2015), could also be broadly effective in general for tumours with aberrant mitochondrial activity.

This study provides comprehensive evidence that both the frequency and types of mutations in genes in cancer are strongly influenced by a given gene’s evolutionary age and its regulatory functions. This furthers our understanding of how a limited number of genetic alterations could promote rapid tumour development through loss of multicellular features and provides an explanation as to how the widespread convergence to common hallmark phenotypes in solid cancer may occur. As we show, this approach can be used to prioritize genes as drivers and identify possible targeted therapies, creating a novel analytical framework that will become increasingly informative as the volume and resolution of cancer genomics data continue to increase.

## Acknowledgements

This work was supported by a Melbourne International Engagement Award (MIEA) and a Melbourne International Fee Remission Scholarship (MIFRS) from the University of Melbourne to A.S.T and funding from the National Health & Medical Research Council of Australia (NHMRC) (APP1052904) and the Peter MacCallum Cancer Foundation to D.L.G., NHMRC Senior Research Fellowships to A.T.P, and a National Health and Medical Research Council (NHMRC) of Australia Program Grant (#1053792) and Fellowship to R.B.P.

The authors declare no competing interests.

## Methods

### Gene ages

The evolutionary ages of genes were obtained from previously published work (Trigos et al., 2017). Phylostratigraphy (Domazet-Loso, Brajkovic, & Tautz, 2007) was used to map human genes onto a phylogenetic tree of 16 phylostrata, spanning genes found across all organisms (Phylostratum 1) to those specific to humans (Phylostratum 16). Human genes with orthologs in primitive unicellular species were assigned to older phylostrata (phylostrata 1-3) and referred to as unicellular (UC) genes, those with orthologs in early metazoan species (phylostrata 4-9) are referred to as early metazoan (EM) genes, and those assigned to phylostrata 10-16 are considered to be mammal-specific (MM) genes.

### Point mutation data

We obtained somatic point mutation data from the Genomic Data Commons Data Portal (https://portal.gdc.cancer.gov/) for 7 tumour types: lung adenocarcinoma (LUAD), lung squamous-cell carcinoma (LUSC), breast adenocarcinoma (BRCA), prostate adenocarcinoma (PRAD), liver hepatocellular carcinoma (LIHC), colon adenocarcinoma (COAD) and stomach adenocarcinoma (STAD). We selected the intersect of variants called by MuSE (Fan et al., 2016), MuTect2 (Cibulskis et al., 2013), VarScan2 (Koboldt et al., 2012) and SomaticSniper (Larson et al., 2012) by The Cancer Genome Atlas. We excluded variants found in ExAC, 1000 genomes, and only included variants from canonical transcripts not found in the last exon. We also excluded genes with large number of known false positives based on previous studies (M. S. Lawrence et al., 2013), namely titin, mucins, ryanodine receptors, dyneins, PCLO, cub- and sushi-domain proteins, neurexins, contactins, PARK2 and olfactory receptors.

Point mutations were classified as a synonymous mutation, missense mutation (missense variant, inframe deletion or inframe insertion), or loss-of-function (LoF) mutation (frameshift, stop-gain, splice acceptor variant and splice donor variant).

### Copy-number aberration data

We obtained copy-number aberration (CNA) data from the Genomic Data Commons Data Portal (https://portal.gdc.cancer.gov/) obtained with SNP arrays for 7 tumour types: LUAD, LUSC, BRCA, PRAD, LIHC, COAD and STAD. Only chromosome regions with at least 10 probes were considered. GenomicRanges (M. Lawrence et al., 2013) was used to assign chromosome regions to genes. Those with positive segment means were considered to be amplifications, and the rest deletions. A gene was considered amplified or deleted only in cases where genes were covered entirely by a region with a CNA. Amplifications were assigned the maximum segment mean and deletions the minimum.

### Enrichment of recurrent point mutations and CNAs in phylostrata

We defined a gene as having recurrent point mutations if there were at least 3 patients across the cohort with missense or LoF mutations in this gene. To account for the background mutation rate of the gene, only genes with a larger number of patients with missense or LoF mutations than with synonymous mutations were considered. We selected recurrent CNAs as those that were amplified or deleted genes in at least 10% of the patients. Note that these procedures were followed for each tumour type independently.

We calculated the fraction of CNA or point mutated genes in each phylostratum for each tumour type as follows:

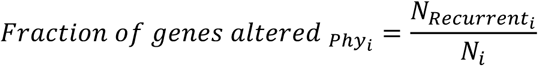

Where *N_Recurrent__i_* corresponds to the number of genes with a recurrent genetic alteration in phylostratum *i*, and *N_i_* the total number of genes in phylostratum *i*.

To account for differences in ranges between tumour types, we ranked the phylostrata by the fraction of genes altered, ranging from 1 (most altered) to 16 (least altered).

To compare the trends of recurrent with non-recurrent alterations, we calculated the fraction of altered genes in each phylostratum that were non-recurrent.

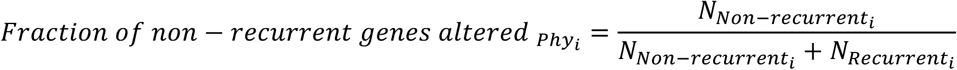

Where *N_Non-recurrent__i_* corresponds to the number of genes with a non-recurrent genetic alteration in phylostratum *i*, and *N_Recurrent__i_* the number of genes with recurrent alterations in phylostratum *i*.

### Gene expression analysis of mutated genes

RNAseq gene expression data from the 7 tumour types and the corresponding normal samples were obtained from The Cancer Genome Atlas.

To determine how the mutational status of a gene affected its expression, we compared the expression of each gene with missense mutations in at least 3 patients or LoF mutations in at least 2 patients in the tumour type cohort against their levels in patients where the gene was unmodified. One-sided Wilcoxon tests were used to determine whether a gene was significantly over or underexpressed in patients where the gene was mutated, using a p-value cut-off of 0.05. We subsequently calculated the ratio of the number of up- and downregulated genes in each phylostratum.

We performed a similar procedure to calculate the effect of point mutations in regulators over the expression of their downstream targets, comparing the level of expression of each downstream target in tumour samples with and without the regulator being mutated using two-sided Wilcoxon tests (p < 0.05). In this case, however, we pooled samples across tumour types to increase power due to the small number of point mutations that occur in regulators. Specifically, we only considered regulators that were point mutated in at least 3 samples across tumour types, and compared the expression of their downstream targets against those of samples of the tumour types where the regulator was not mutated.

The magnitude of the downstream effect of mutating regulators was classified as low (<5% down downstream targets affected), moderate (5-20%) or high (>20%) impact. We defined the prevalence of EM genes as regulators as the ratio of the number of EM regulators over the number of UC regulators. The log10 values were calculated to normalize to zero.

### Signatures of selection in chromosomes

We defined the fraction of copy-number aberrant chromosome for each patient and chromosome as the ratio of the number of genes affected by amplifications or deletions and the total number of genes in the chromosome. To associate this chromosomal context with evolutionary ages of genes, we determined the presence or absence of genes of a specific phylostratum in the copy-number aberrant chromosome regions of each patient. The information was aggregated across patients by averaging the fraction of chromosome altered for each chromosome and phylostratum. Genes in shorter CNA regions (smaller fraction) were considered to be under stronger, focal selection, whereas CNAs in genes found in longer CNA regions (larger fraction) were considered to result from broad amplifications or deletions.

We defined focally amplified or deleted genes as those in which their chromosomal context was in the upper quantile (0.25) of the distribution of the mean fraction of copy-number aberrant chromosome across patients for each chromosome.

### Gene regulatory network analyses

We obtained a human gene regulatory network (GRN) from PathwayCommons (version 9) by selecting pairs of genes connected by an edge of the type “control-expression-of”, resulting in a directed network with 95,651 edges. We defined regulator genes as those with at least one downstream target.

We defined ‘out-degree’ as the number of outgoing edges of a regulator, representing the extent of its downstream regulatory network. In contrast, the ‘in-degree’ of a gene was defined as the number of incoming edges and it is proportional to how highly regulated the gene is. Greater out-degree/in-degree ratios indicate bias towards a higher number of outgoing edges (i.e. regulatory role), whereas smaller ratios indicate bias towards a higher number of incoming edges (i.e. target role).

We also obtained protein-protein (PPI) networks of humans from PathwayCommons (Cerami et al., 2011) version 9, BioGRID (Chatr-Aryamontri et al., 2017) version 3.4.152 and the InWeb_IM network (Li et al., 2017). Only nodes and edges corresponding to genes and links between two genes were considered. Since these networks are undirected, we only calculated the degree of a gene as the total number of edges associated with the gene.

### Classification of regulators by the evolutionary ages of their downstream targets

Regulators were classified as UC-t, EM-t, MM-t or UC/EM-i regulators based on the ages of their target genes, independently of the evolutionary age of the regulator. First, we calculated the percentage of downstream UC, EM and MM target genes for each regulator. A regulator was classified as UC-t if more than 2/3 of its targets were UC genes, as EM-t if more than 2/3 of its targets were EM genes, or MM-t if more than 2/3 if its targets were MM. Regulators that did not meet the above criteria, but at least 1/10 of their target genes were UC genes or EM genes, and less than 1/10 were MM genes, were classified as UC/EM-i regulators.

### CNA in regulators and targets

To determine whether the copy-number status of a regulator was associated with the fraction of downstream CNA targets, we calculated the fraction of downstream CNA targets for each regulator in each patient. We compared the fraction of CNA downstream target genes when regulators were CNA and CNN for each patient, and used Wilcoxon tests to determine if there were significant differences in these fractions between patients where the regulator was CNA or CNN. We only considered regulators that were CNA and CNN in at least 3 patients across all tumour cohorts, and had at least 2 targets. P-values were corrected for multiple testing using Benjamini-Hochberg correction.

A summary fraction of CNA targets was obtained for each regulator by calculating the median across patients, and we calculated the difference in percentage of targets with CNAs by subtracting the fraction of targets when the regulator was CNA minus the fraction when the regulator was CNN.

### Gene expression analyses of CNA targets and regulators

RNAseq gene expression data from the 7 tumour types and the corresponding normal samples were obtained from The Cancer Genome Atlas. We evaluated the effect of CNAs on gene expression by calculating the expression fold-change between matched tumour and normal samples for each gene, and comparing the fold-changes of samples where the gene was CNA and where it was CNN using one-sided Wilcoxon tests. Only genes that were amplified or deleted in at least 3 samples were considered. Benjamini-Hochberg correction was used for correction for multiple testing. We next calculated the percentage of significantly over or underexpressed amplified or deleted UC, EM and MM target genes, respectively. We calculated the percentage of differentially expressed CNA targets as the ratio between the number of differentially expressed CNA targets of each regulator and the total number of CNA target genes. We subsequently calculated the median percentage of differentially expressed CNA targets for each regulator class in each tumour type. Wilcoxon tests were calculated on the median values per regulator across tumours to determine significant differences.

### Cancer cell line gene knockout, mutation and IC50 values

Scores of the probability of dependency to genes across 364 solid-tumour tissue cell lines were obtained from the Avana CRISPR-Cas9 genome-scale knockout dataset generated by Project Achilles and the Cancer Dependency Map project (18Q1 version) (https://portals.broadinstitute.org/achilles). We excluded all cell lines from haematopoietic and lymphoid tissues. A cell line was considered to be dependent on a regulator if its probability of dependency was greater than 0.95. The enrichment of a regulator class among the regulators to which cancer cell lines were dependent was determined using the odds ratio.

Mutation and CNA information of cell lines was obtained from the Cancer Cell Line Encyclopaedia (CCLE) (https://portals.broadinstitute.org/ccle). Significant differences in the dependency scores of cell lines with mutated and non-mutated regulators, or amplified or copy-number normal regulators were obtained using Wilcoxon tests (p < 0.05). We only considered regulators that are either amplified or point mutated in at least 3 cell lines.

IC50 values of cancer cell lines after treatment with 250 cancer drugs were obtained from the Genomics of Drug Sensitivity in Cancer database (version 17) (Yang et al., 2013). We calculated the Spearman correlation of IC50 values with dependency scores from the Avana CRISPR-Cas9 databases. We only considered negative correlations < −0.25 and with a adjusted p-values after Benjamini and Hochberg correction < 0.05, since we were interested in cell lines with high dependency to a regulator and that showed greater drug sensitivity at lower concentrations.

